# Transcriptional pathobiology and multi-omics predictors for Parkinson’s disease

**DOI:** 10.1101/2024.06.18.599639

**Authors:** Ruifeng Hu, Ruoxuan Wang, Jie Yuan, Zechuan Lin, Elizabeth Hutchins, Barry Landin, Zhixiang Liao, Ganqiang Liu, Clemens R. Scherzer, Xianjun Dong

**Affiliations:** APDA Center for Advanced Parkinson Research, Brigham and Women’s Hospital, Harvard Medical School, Boston, MA, USA; Precision Neurology Program, Brigham & Women’s Hospital, Harvard Medical School, Boston, MA, USA; Genomics and Bioinformatics Hub, Department of Neurology, Brigham and Women’s Hospital, Harvard Medical School, Boston, MA, USA; Aligning Science Across Parkinson’s (ASAP) Collaborative Research Network, Chevy Chase, MD, USA; Neurogenomics Division, Translational Genomics Research Institute (TGen), Phoenix, AZ, USA; Technome, Herndon, Virginia, USA; Shenzhen Key Laboratory of Systems Medicine in Inflammatory Diseases, School of Medicine, Shenzhen Campus of Sun Yat-sen University, Shenzhen, Guangdong, China

**Keywords:** Parkinson’s disease, Differentially expressed genes, Neutrophil activation, Machine learning

## Abstract

Early diagnosis and biomarker discovery to bolster the therapeutic pipeline for Parkinson’s disease (PD) are urgently needed. In this study, we leverage the large-scale whole-blood total RNA-seq dataset from the Accelerating Medicine Partnership in Parkinson’s Disease (AMP PD) program to identify PD-associated RNAs, including both known genes and novel circular RNAs (circRNA) and enhancer RNAs (eRNAs). There were 1,111 significant marker RNAs, including 491 genes, 599 eRNAs, and 21 circRNAs, that were first discovered in the PPMI cohort (FDR < 0.05) and confirmed in the PDBP/BioFIND cohorts (nominal *p* < 0.05). Functional enrichment analysis showed that the PD-associated genes are involved in neutrophil activation and degranulation, as well as the TNF-alpha signaling pathway. We further compare the PD-associated genes in blood with those in post-mortem brain dopamine neurons in our BRAINcode cohort. 44 genes show significant changes with the same direction in both PD brain neurons and PD blood, including neuroinflammation-associated genes *IKBIP*, *CXCR2*, and *NFKBIB*. Finally, we built a novel multi-omics machine learning model to predict PD diagnosis with high performance (AUC = 0.89), which was superior to previous studies and might aid the decision-making for PD diagnosis in clinical practice. In summary, this study delineates a wide spectrum of the known and novel RNAs linked to PD and are detectable in circulating blood cells in a harmonized, large-scale dataset. It provides a generally useful computational framework for further biomarker development and early disease prediction.

**Significance statement:** Early and accurate diagnosis of Parkinson’s disease (PD) is urgently needed. However, biomarkers for early detection of PD are still lacking. Also, the limit of sample size remains one of the main pitfalls of current PD biomarker studies. We employed an analysis of large-scale whole-blood RNA-seq data. By identifying 1,111 significant marker RNAs, we establish a robust foundation for early PD detection, which implicated in neutrophil activation, degranulation, and TNF-alpha signaling, offer unprecedented insights into PD pathogenesis. Our multi-omics machine learning model, boasting an AUC of 0.89, outperforms previous studies, promising a transformative tool for precise PD diagnosis in clinical settings. This study marks a pivotal step toward enhanced biomarker development and early disease prediction.

## Introduction

Parkinson’s disease is a progressive neurodegenerative disease, which causes the loss of dopamine neuron cells in the substantia nigra pars compacta of the human midbrain. It is estimated that the number of PD patients is expected to exceed 13 million by 2040 globally(1, 2). PD is thought to be caused by combinatorial effects of environmental, epigenetic, and genetic contributions that exert many of their effects through cis- and trans-acting regulation of transcript abundance(3–6). Progressive loss of dopamine neurons and an increasing burden of α-synuclein-positive neuronal inclusions (the so-called Lewy bodies) are hallmarks of PD(7, 8), once PD neuropathology crosses a clinically relevant threshold, movement becomes relentlessly more impaired in PD patients.

Biomarkers for early detection and quantitative tracking of disease progression are currently lacking(9, 10). By the time a patient is diagnosed with PD based on today’s clinical criteria (e.g. resting tremor, slow movements, and stiffness), ∼70% of vulnerable dopaminergic neurons have been lost(11). Therefore, developing a panel for early and accurate diagnosis is urgently needed(12, 13).

Additionally, PD is a slowly progressive, complex genetic disease that likely results from multiple genetic risk variants, each conferring small increases in susceptibility. PD GWAS have revealed thousands of genetic variants whose mutations are associated with disease risk(14). However, they are static in the germline and thus cannot be used to quantitatively track disease progression over time in serial measures. While the strategy of constructing an aggregate measure from multiple individual markers has been fruitful in genetic studies of PD risk, the use of markers spanning multiple modalities (*e.g.,* genetic, transcriptomic, clinical, and imaging-based markers) is needed to maximize its utility(15), as it is unlikely that a single biomarker will adequately capture this genetic and environmental heterogeneity. Also, individuals whose risk of developing PD is high may show developmental potentials in their transcriptomics profiles before having the clinical detective symptoms, even if they may pass the clinical PD tests(16, 17). Limited sample size remains one of the main pitfalls in current biomarker studies. A systematic review of published studies using α-synuclein species as a PD biomarker found that 84% of studies included 100 PD patients or fewer(18). Previous efforts have made several individual cohorts available to study PD biomarkers, including the Michael J. Fox Foundation (MJFF) Parkinson’s Progression Markers Initiative (PPMI)(19, 20), the NINDS Parkinson’s Disease Biomarkers Program (PDBP)(21), and the MJFF BioFIND study(22). All these data have been integrated into the Accelerating Medicine Partnership in Parkinson’s Disease (AMP PD) program, which to date has generated longitudinal RNA sequencing data for about 8,500 samples covering more than 3,200 participants with deep clinical phenotype data. A study by Craig *et.al.*(23) conducted an overview analysis of the PPMI dataset, finding that the neutrophil cell abundance is higher in patients with PD, while lymphocyte cell abundance is lower in patients with PD. However, they were using all time-point samples and treated each sample at a time-point as an independent individual, the results may not be representative for early time-points samples. In another study by Makarious *et.al.,* the authors built multi-modal machine learning models, which have good performances, however, the case/control samples are unbalanced, and they developed models just achieving modest balanced accuracy(13).

In our work, we leveraged the large-scale RNA-seq dataset in the AMP PD baseline to find the most significant differentially expressed genes (DEGs, which were mainly known mRNAs or lncRNAs) and novel non-coding RNAs which include circular RNAs (circRNAs) and enhancer RNAs (eRNAs), between PD patients and healthy controls in a defined discovery dataset. We validated these PD-associated RNAs in an independent replication cohort. Enrichment analysis was conducted to find the distinguishable functional pathways or gene ontology terms between PD patients and healthy controls using the replicated differentially expressed protein-coding genes. We then built the multi-gene classifiers using a selective set of PD-associated marker genes, as well as an integrative multi-omics classifier using gene expression data, genetic variant data, and clinical information to achieve better performance, in our study, all models are trained on the discovery dataset and tested on the replication dataset to evaluate their respective performance. We developed and tested innovative multi-omics classifiers that would provide reusable computational framework for PD diagnosis in clinical practices.

## Results

### Data sources

This study used data from the AMP PD program, which includes eight cohorts, including PPMI, PDBP, and BioFIND) with RNA-seq data for 3,274 participants. Whole genome sequences (WGS) and clinical data are also available. The PPMI is a longitudinal observational study of 1,923 participants with PD or at risk for PD and healthy volunteers, contributing comprehensive clinical and imaging data and biological samples at 33 clinical sites around the world. In the PPMI study, Patients in the PD cohort have been diagnosed within 2 years of enrollment. Importantly, they are de novo patients and as such not yet taking PD medications, which may confound biomarker analyses. The PDBP is another PD program that collect standardized longitudinal clinical data and biospecimens across all stages of PD, which has 1,604 participants and was developed to accelerate the discovery of promising new diagnostic and progression biomarkers for PD. The PDBP supports basic, translational and clinical research through hypothesis testing, target and pathway discovery, biomarker development, and disease modeling. BioFIND is an observational clinical study designed to discover and verify biomarkers of PD, that includes blood and cerebrospinal fluid, in 118 well-defined, moderately advanced people with PD and 88 control volunteers. We obtained permission to access the AMP PD datasets on Google Cloud Storage for the analysis. All information on data collection and data processing procedures from the AMP PD program can be found at https://www.amp-pd.org.

### Summary of data quality

In this study, to systematically delineate genes associated with PD diagnosis at an early stage, we leveraged primarily the baseline data (visit month=0). The PPMI cohort was used as discovery population. PDBP/BioFIND cohorts were defined as the replication population (**Fig 1**). Most of the PD cases in the replication dataset are moderate and advanced, while most PD cases in PPMI are recruited when they are newly diagnosed ((**Fig S1**). We intend to focus on individuals recently diagnosed with PD, pinpointing early stage DEGs. Subsequently, we plan to validate these DEGs using samples from advanced PD to establish a set of DEGs exhibiting stable changes. These genes may serve as diagnosis markers at an early stage. We applied stringent quality check standards to the samples and to the genes (see Methods for details). After the stepped sample filtrations, there were 551 case samples, and 437 control samples were kept in the discovery dataset (**Fig S2**). For the replication dataset, 760 case samples and 452 control samples were used for the following analysis (**Fig S3**). After removing low abundance and low variance genes, 35,900 genes were available in the discovery dataset. The sex check shows high consistency between clinically reported sex and genetically determined sex (**Fig S4A**). The case/control sample distributions on the plates show that the percentages of healthy samples on some plates (P201, P207, P208, P209, P210, P211, P212, P213, P214, and P215) are higher than others in the replication dataset, but there is no extreme plate outlier (**Fig S4B**). Therefore, we included the plate as an effect factor in our DEseq2 formula design. Based on our data quality assessments, there are no extreme outliers, so we did not remove any extra samples. For the replication dataset, 31,935 genes were kept. The scatter plot demonstrated the sex check consistency between clinically reported sex and genetically determined sex (**Fig S4C**). And the case/control sample distributions on the plates show more even distributions across all plates (**Fig S4D**) Finally, 988 baseline samples were included in the discovery dataset and 1,212 baseline samples were included in the replication dataset for all the downstream analyses. The basic clinical characteristics of the datasets after filtrations are summarized in **Table 1**.

**Fig 1.**
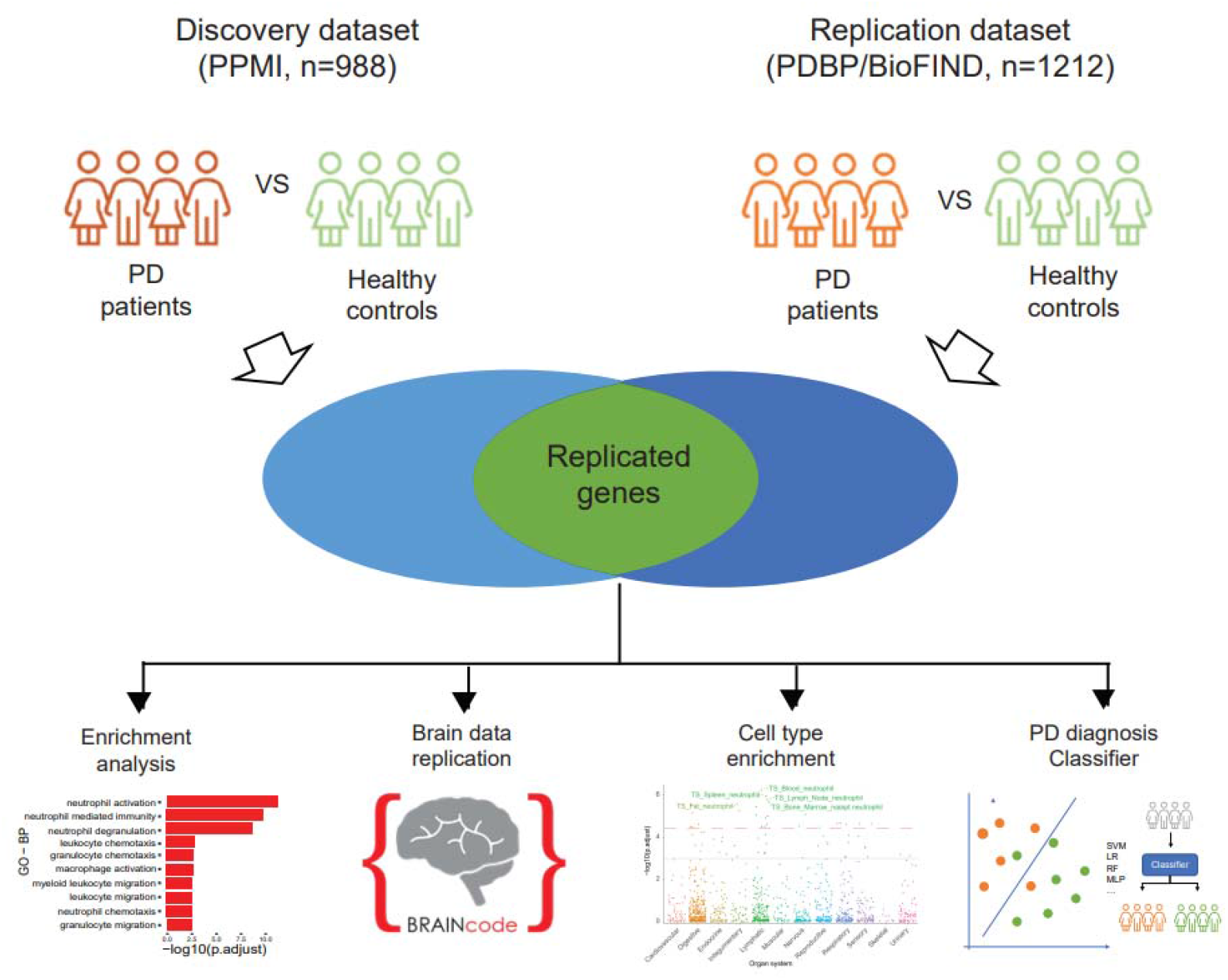
The general workflow of our study. The study was designed in cross-sectional approach. Significant DEGs/DE eRNAs/DE circRNAs were discovered from PPMI cohorts and replicated in PDBP/BioFIND cohort. Further analysis such as functional enrichment analysis, replication with brain sample data and cell type enrichment analysis were conducted on the replicated DEG. The PD diagnosis classifiers were built and tested utilizing the multi-omics data.

**Table 1.**
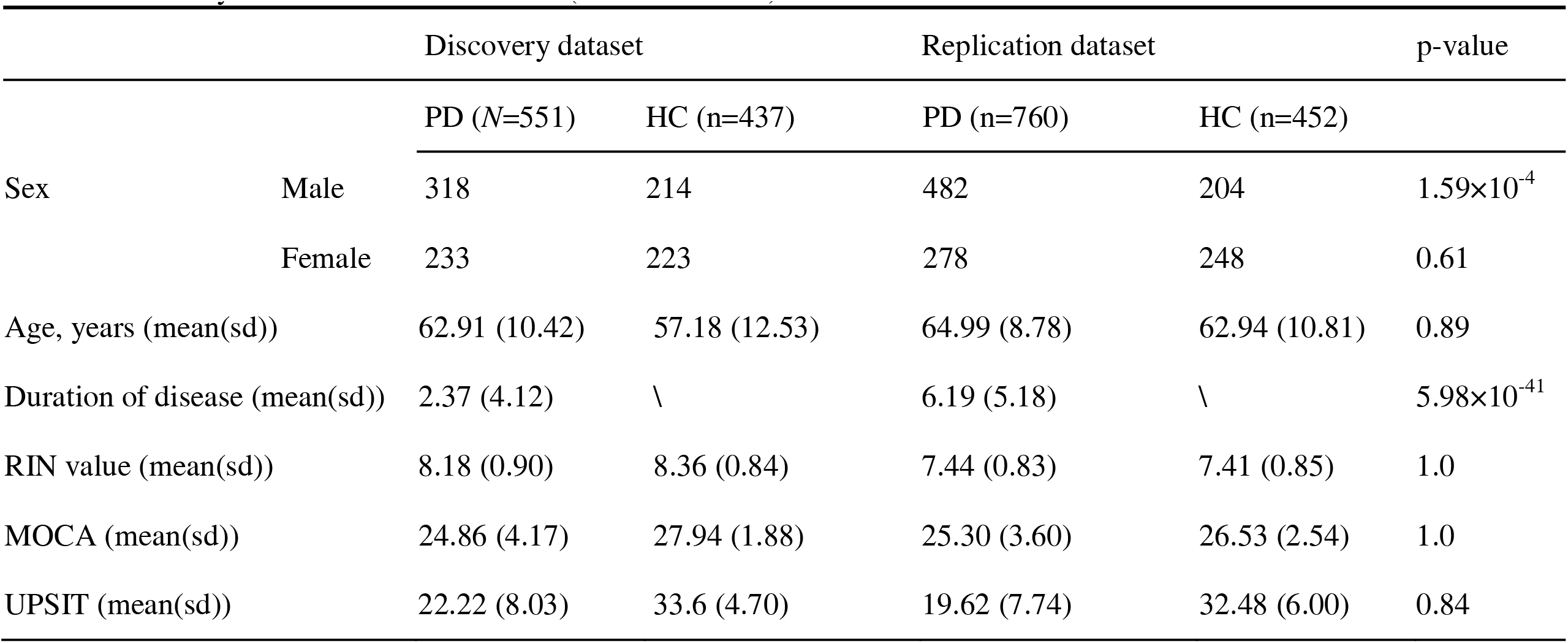
Summary of clinical data at baseline (after filtrations.)

### Gene expression associated with PD

Through the differential expression analysis of the genes in the discovery dataset, 874 DEGs were discovered in the discovery dataset with Benjamin-Hochberg adjusted p-value < 0.05 (**Fig 2**). Of these, 502 genes were replicated in the replication dataset with a nominal p-value < 0.05 (**Fig 2**), of which over 97.8% (491) genes have consistent direction-changes in both discovery and replication datasets (**Fig 2**, and the details are available in **Supplementary Table S1**). The over-representation enrichment analysis was conducted on GO and WikiPathway terms using 491 replicated DEGs with the same change directions. There are 5 significantly enriched GO biological processes (GO-BP) with FDR < 0.05, as well as 5 significantly enriched GO cellular component (GO-CC) terms (**Figs 3A**, **Supplementary Table S2**). The results show that biological processes of neutrophil activation and degranulation were significantly enriched with the lowest adjusted p-values of 4.64*10^-5^. For the enriched GO-CCs, most are granule and membrane-related terms. Both the enriched GO biological processes and cellular components revealed that neutrophil activation and neutrophil degranulation are the key messages derived from the DEGs. Additionally, in the enriched GO-BP terms, we found that the genes are also involved in immune-related pathways, which was also confirmed by the enriched WikiPathways results (**Fig 3B**, **Supplementary Table S2**).

**Fig 2.**
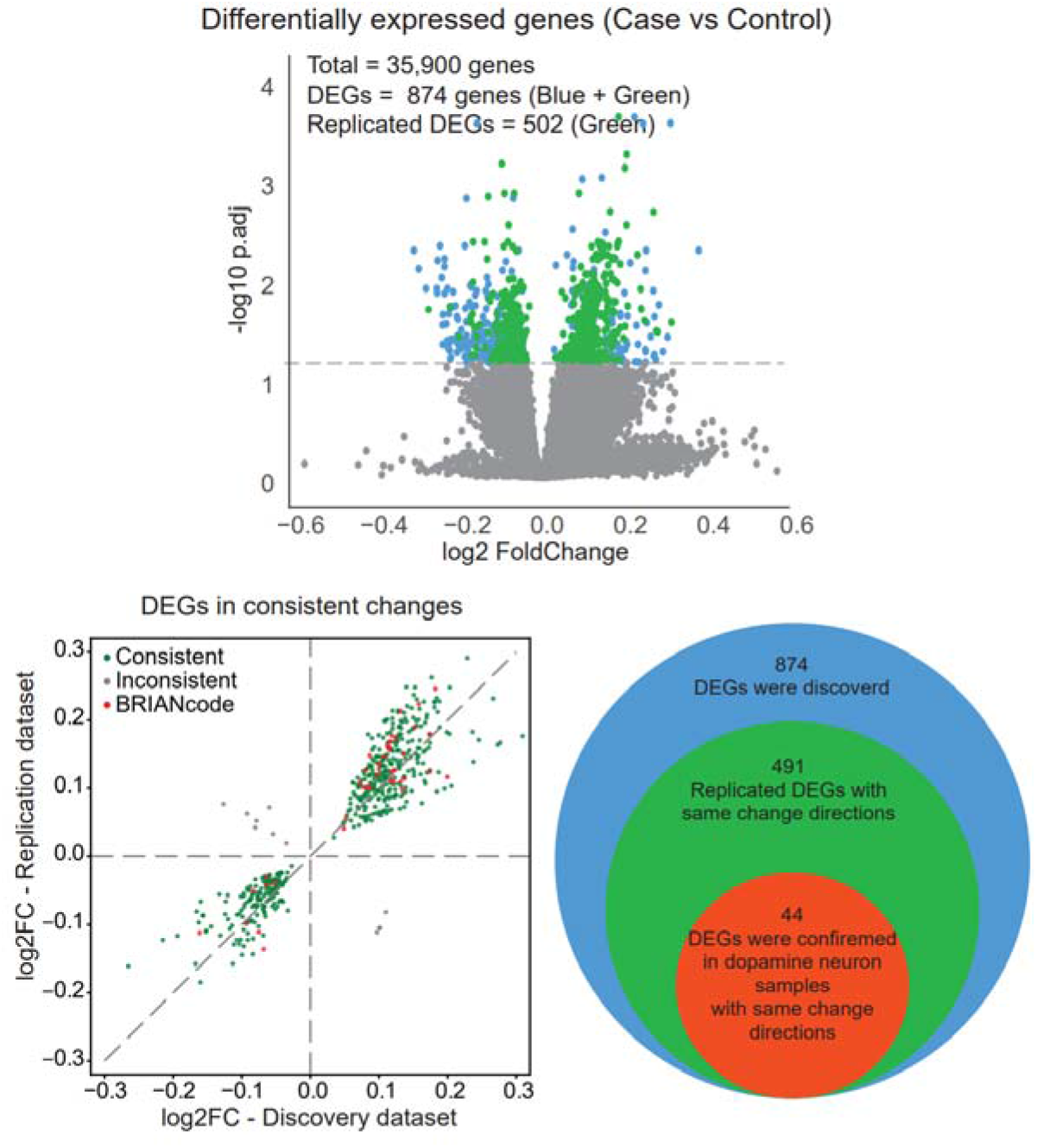
Results of differentially expressed gene analysis. The volcano plot shows that 874 DEG were discovered from the discovery dataset with FDR < 0.05 (the blue and green dots). Then 502 genes were replicated with a nominal p-value < 0.05 in the replication dataset (the green dots). The scatter plot and the Venn diagram show among the 502 replicated genes, 491 replicated DEGs have consistent change directions in both discovery and replication datasets. Forty-four replicated DEGs with consistent change directions were further confirmed in dopamine neuron samples.

**Fig 3.**
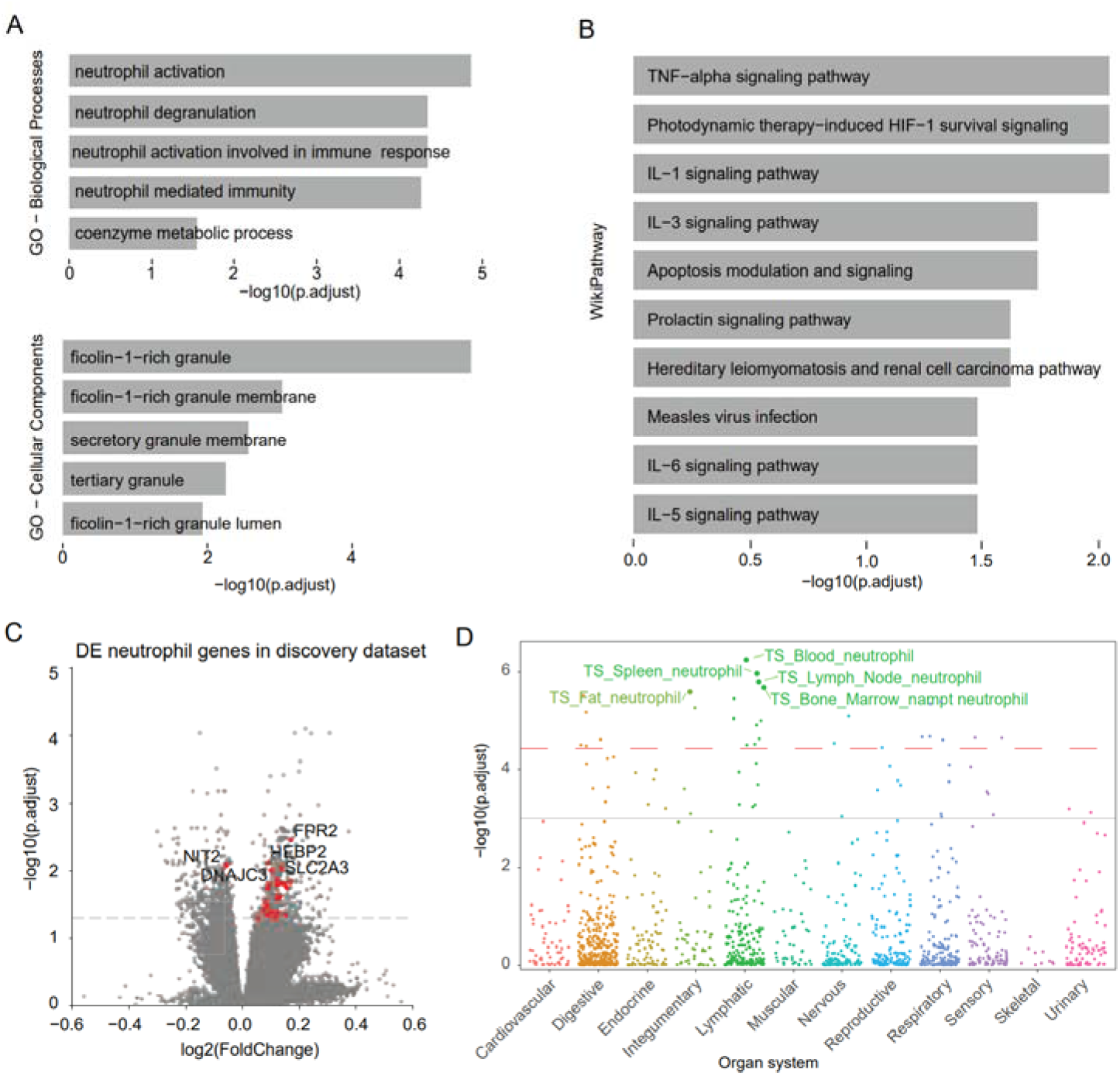
The enrichment analysis results of the replicated differentially expressed genes and the replicated differentially expressed neutrophil genes. (A) The enriched GO-BO and GO-CC terms with adjust p < 0.05, the neutrophil degranulation and neutrophil activation are significantly enriched. And the ficolin-1-rich granule and membrane cell components were significantly enriched. (B) Both the enriched GO-BP and the WikiPathway show that DEGs are involved in immune-related pathways. (C) The volcano plot of replicated differentially expressed neutrophil genes in discovery datasets shows that most neutrophil genes are up-regulated in PD patients (the top 5 ranked genes were shown on the plot, the full list can be found in **supplementary Table S2**). (D) Cell-type specific enrichment analysis shows the replicated genes are enriched in neutrophil cell types in the lymphatic system.

By looking into the changes of the leading genes enriched in the neutrophil activation and neutrophil degranulation biological processes, we found that 29 DEGs are involved in these two GO-BPs and of which 28 genes were upregulated in PD case samples in both the discovery and the replication datasets (**Fig 3C**, **Supplementary Table S2**). The results indicate that the neutrophil activation and the neutrophil degranulation were highly regulated in PD patients. Furthermore, the highly expressed neutrophil genes or the highly regulated neutrophil activation and neutrophil degranulation can serve as the biomarkers for PD early diagnosis.

In the analysis of cell-type-specific enrichment, the 491 DEGs were also highly enriched in the blood neutrophil cell of the lymphatic organ system (**Fig 3D**, the dashed red line in the plot indicates the significant threshold (p = 3.69 x 10^-5^) corrected with 1355 collected tissue-cell types. The solid grey line indicates the nominal significance (p = 0.001)). This further confirms that the upregulation of neutrophil genes could be a markers of early PD diagnosis.

Among these 29 differentially expressed leading genes in neutrophil activation and neutrophil degranulation biological processes, several have been studied in the context of PD. A pathogenic mutation (p.N855S) in *DNAJC13* was linked to autosomal dominant Lewy body PD(24–26). *Apaf-1*(apoptotic peptidase activating factor) was reported as a potential drug target for neurodegenerative diseases and *Apaf-1* dominant negative inhibitor can prevent *MPTP* toxicity as antiapoptotic gene therapy for Parkinson’s disease(27). *FCGR2A* and *FCGR2B* are involved in phagocytosis and modulate inflammatory responses and mutated *FCGR2A* is causal to Parkinson’s disease(28). *FCGR2B* expressed in neurons functions as a receptor for a-syn fibrils and mediates cell-to-cell transmission of a-synuclein and the *FCGR2B-SHP-1/-2* signaling pathway may be a therapeutic target for the progression of PD(29). The *CD93* participates in pathophysiological processes of central nervous system inflammation(30).

*SNCA* is an inevitable gene when it comes to PD. We validated the reduced SNCA expression trends in PD samples than health controls in multiple cohorts (including PPMI, PDBP/BioFIND, BRAINcode) and on multiple platforms (including RNAseq and NanoString) (**Fig 4**). Our findings showed the consistent lower *SNCA* mRNA levels in blood of early-stage PD patients with the brain samples, which also align with the Locascio’s findings in HBS cohort (31, 32).

**Fig 4.**
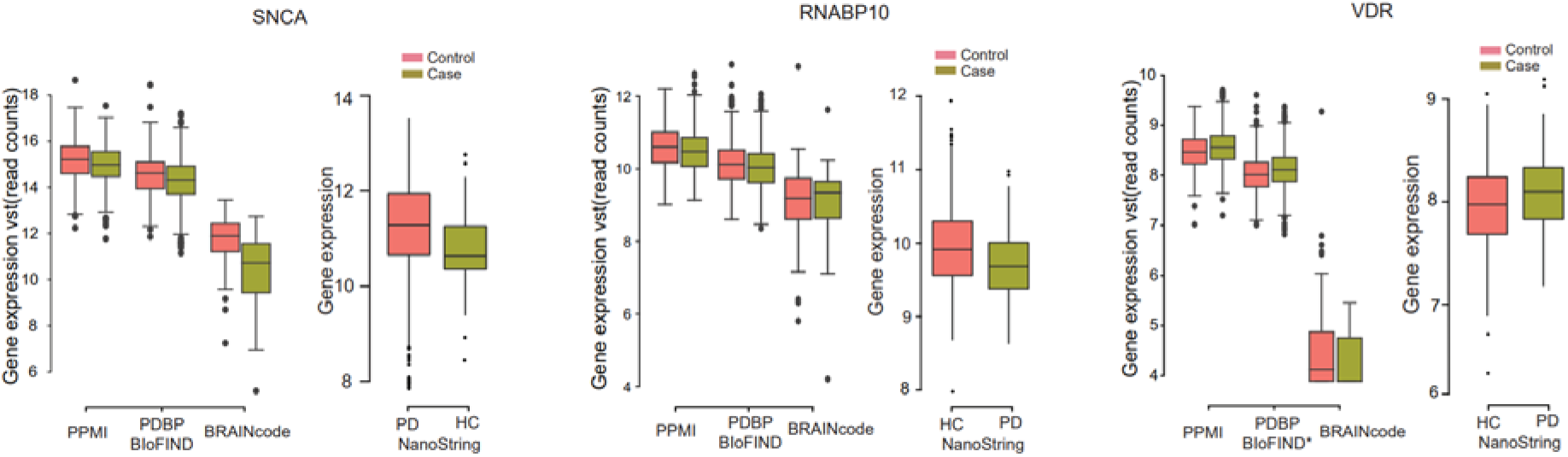
The expression levels of three PD-associated genes *SNCA, RNABP10, and VDR* in the discovery, replication, BRAINcode datasets, and NanoString dataset.

We also replicated pathological levels of *VDR* and *RANBP10*(33). Vitamin D is linked to neuroprotection in animal models of PD(34) and we previously reported reduced levels of the vitamin D receptor (*VDR*) in an unbiased microarray screen of PD blood samples (33) and 25-hydroxy-vitamin D deficiency in 17.6% of PD patients(34). For these previously nominated candidate biomarker RNAs of PD, the same direction-change were observed between PD and health control samples in both the NanoString data and RNA-seq data (PPMI, PDBP/BioFIND) (**Fig 4**).

Our secondary analysis tested the expression changes associated with motor severity which were indicated by the MDS-UPDRS part III summary score. Our results are shown in the **Supplementary Table S3**. We found that 2,236 genes and 4,045 genes were significantly associated (adjust p < 0.05) with the MDS-UPDRS part III summary scores in the discovery and replication datasets respectively. Among these genes, 1,636 genes were shared by both the discovery and the replication datasets with the same change directions. GO enrichment analysis was conducted on the 1,636 replicated genes, and the results showed that the “neutrophil activation”, “neutrophil activation involved in immune response”, “neutrophil-mediated immunity” and “neutrophil degranulation” are the top enriched GO-BPs. This is consistent with our conclusion from the main DE analysis between the PD cases and healthy controls. These findings suggest that neutrophil degranulation are potential biomarkers in blood for PD diagnosis.

### Replicating blood-based marker genes in brain neurons

In our DE analysis of the brain neuron data, we identified 575 DEGs with Benjamin-Hochberg adjusted p-value < 0.05. Compared with the 491 consistently changed blood-DEGs in both discovery and replication, 44 replicated DEGs with constant direction-changes were further confirmed in dopamine neuron samples (**Fig 2, Supplementary Table S1**). The 44 replicated DEGs with constant change directions shared 5 genes with the 29 differentially expressed circulating genes in neutrophil activation and neutrophil degranulation biological processes. Those genes are *PREX1*, *FCGR2A*, *CAB39*, *CXCR2*, *LAMP2*.

Among these 44 DEGs, the *LAMP2* gene has been reported to be differentially affected between the early stages of PD and controls and was also reported to be association with the expression level of *SNCA*(35). Also, there are several neuroinflammation-associated genes were replicated in our brain datasets, such as *IKBIP*(36)*, CXCR2*(37, 38), and *NFKBIB*(39). Additionally, *IL18R1*, a cytokine receptor that belongs to the interleukin 1 receptor family, is significantly increased in PD blood and brain neurons. While the function of this cytokine receptor in PD is not experimentally verified, an increase in interleukin-1beta (*IL-1*β) was previously reported as a potential mediator of microglia activation in the PD rat model(40). These genes were consistently upregulated.

### Differentially expressed novel RNAs

In our study, we identified a total of 26,035 candidate eRNAs using our in-house scripts(6), all of which surpassed the established low expression filtration threshold (only eRNAs with read count > 5 in >10% samples were kept). Out of these, 783 exhibited differential expression, with 599 of these DE eRNAs replicating the same directional changes in the combined PDBP/BioFIND dataset. The dataset revealed that among the replicated DE eRNAs, 396 were up-regulated, and 203 were down-regulated (**Supplementary Table S4**).

Regarding circRNAs, we defined 441,811 circRNAs. After applying a filtration process requiring at least 2 read counts in 10% of samples, 3,052 circRNAs remained and were subjected to DE analysis. This analysis led to the discovery of 35 DE circRNAs, 21 of which were replicated in the PDBP/BioFIND dataset. Among these replicated DE circRNAs, 15 were up-regulated, and 6 were down-regulated (**Supplementary Table S4**).

We then further investigated the host genes of these replicated DE eRNAs and DE circRNAs. 599 DE eRNAs and 21 DE circRNAs were mapped to 306 host genes (289 eRNA host genes, 18 circRNA host genes, 1 shared host gene (ENSG00000159339, *PADI4*)). Although, the host genes were not sharing the same enriched GO terms with DEGs, we noticed several PD-associated genes or genes that are involved in neutrophil activation in the host gene list. *SPI1* is one of the member genes of GO:0042119 (neutrophil activation). It has been reported that *SPIL1* plays a crucial role in regulation of the genes relevant to specialized functions of microglia, therefore dysregulation of *SPIL1* might contribute to the establishment or development of PD due to the accumulation of activated microglia(41–43). *PADI4* is a gene that can positively regulate *TNF-*α and *CCL2* which can lead to the development of neuroinflammation(44, 45). *PADI2* coordinates with *PADI4* to regulate the assembly of the *NLRP3* inflammasome to promote *IL-1*β release. Research also showed that *PADI4* can participate in all aspects of neutrophil extracellular traps (NETs)(46). Moreover, X-linked dystonia Parkinson’s disease is aggravated by increased levels of *PADI2*, *PADI4*, and inflammation in the prefrontal cortex (PFC) and its derived fibroblasts(47). The circRNA host gene *RHBDD1*, also named *RHBDL4*, has been implicated in a variety of diseases including AD and Parkinson disease, which can cleave amyloid precursor protein inside the cell, causing it to bypass amyloidogenic processing, leading to reduced Aβ levels(48), this gene had a significant negative log2 fold change in PD patients comparing to the health control in both discovery and replication cohorts. Among the 306 host genes, 53 genes were shared with replicated DEGs. The DEGs *IKBIP, LAMP2, VDR* which are associated with PD as we mentioned above were also among the host genes.

### Performance of PD diagnosis classifier models

The PPMI samples (discovery population) were randomly split into a training set (80%) and a validation set (20%) for 100 times, so that we had 100 pairs of training-validation datasets. The training set was used to build the classifiers which were validated on the corresponding validation dataset. All the trained models were tested on the independent replication cohorts. There are 10 different machine learning classifiers were trained and compared (see Methods for details). Since there are too many features, to avoid the overfitting problems, during the model training we used the LASSO for feature selection. The PD diagnosis classifiers were first constructed using the 874 DEGs from PPMI with FDR < 0.05. After feature selection, there are 23 to 36 DEGs that were selected as the final predictors from each random split for model training and validation. Among all the 10 tested algorithms, we observed that the support vector machine with rbf kernel (SVM_rbf) had relatively better performances than others regarding to the average AUROC and the area under the precision-recall curve (AUPRC) values on the PDBP/BioFIND testing dataset (**Fig 5A)**. The mean and standard deviation of AUROC and AUPRC were 0.72 (0.03) and 0.72 (0.04) on the PPMI 20% withheld validation datasets, and 0.64 (0.01) and 0.74 (0.01) when applied the model to the PDBP/BioFIND testing dataset (**Table 2, Supplementary Table S5**). After adding the polygenic risk score (PRS) as the genetics feature to the selected DEGs in each random split, the logistic regression (LR) model demonstrated best performance, the AUROC and AUPRC were improved to 0.75 (0.03) and 0.78 (0.03) on the validation datasets, and 0.70 (0.01) and 0.79 (0.01) on the independent testing dataset (**Fig 5B**, **Table 2**). With the clinical data (UPSIT-smell test score, sex, and age) added, the support vector machine (SVM) emerged as the optimal model. The AUROC and AUPRC values were raised to 0.91(0.02) and 0.92(0.02) on the validation dataset, and 0.89 (0.01) and 0.93 (0.01) on the testing dataset (**Fig 5C**, **Table 2**). **Fig 5D and Fig 5E** show the progressive improvement in model performance with step-wised addition of the genetics and clinical features on the validation and testing dataset.

**Fig 5.**
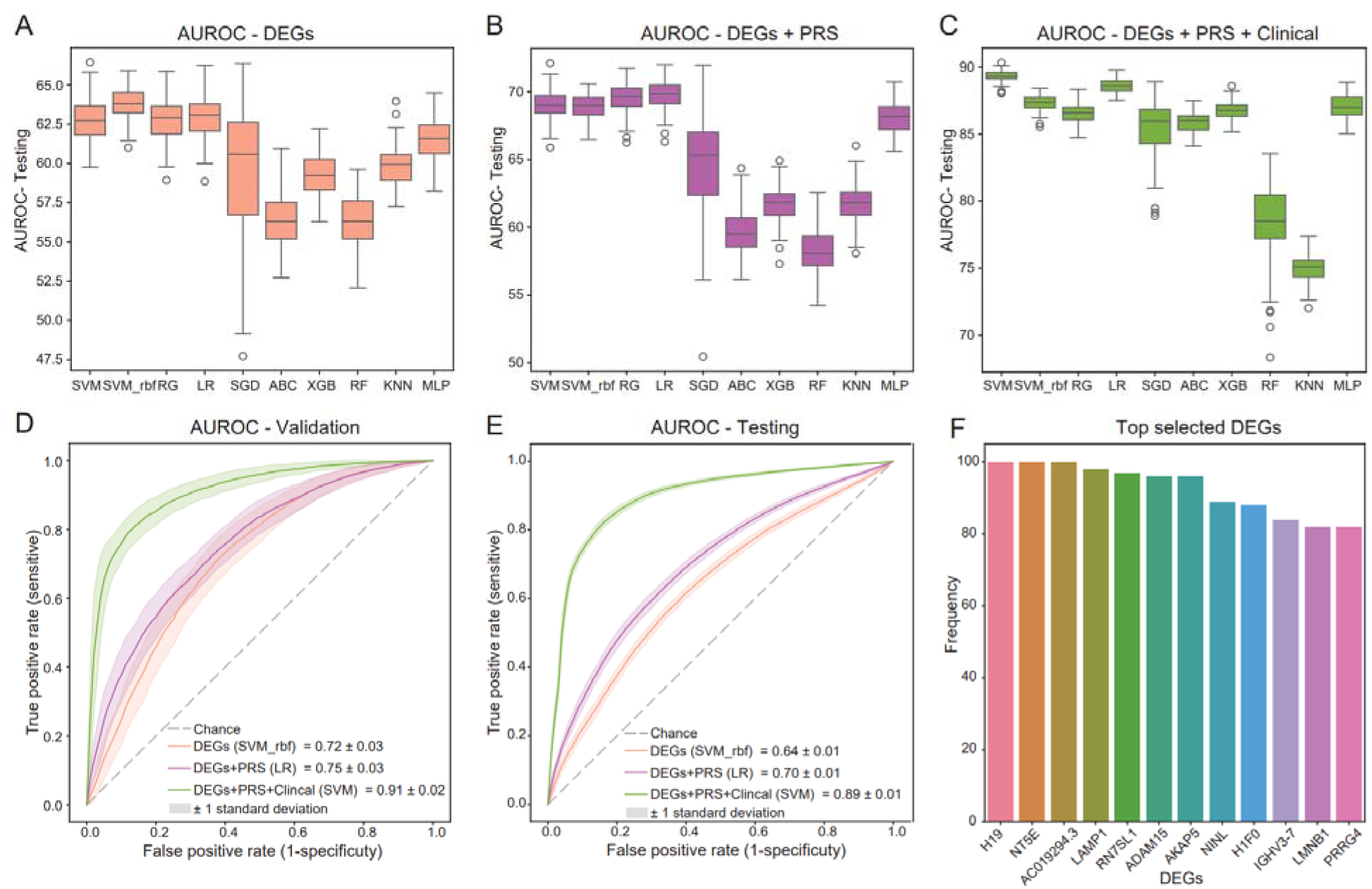
The performance of the models. (A-C) the AUROC value distributions of 10 tested algorithms with step-wised add feature sets on the testing dataset. (D, E) AUROC curves of the best models with step-wised add feature sets on the validation dataset and testing dataset. (F) Top selected genes during feature selection.

**Table 2.**
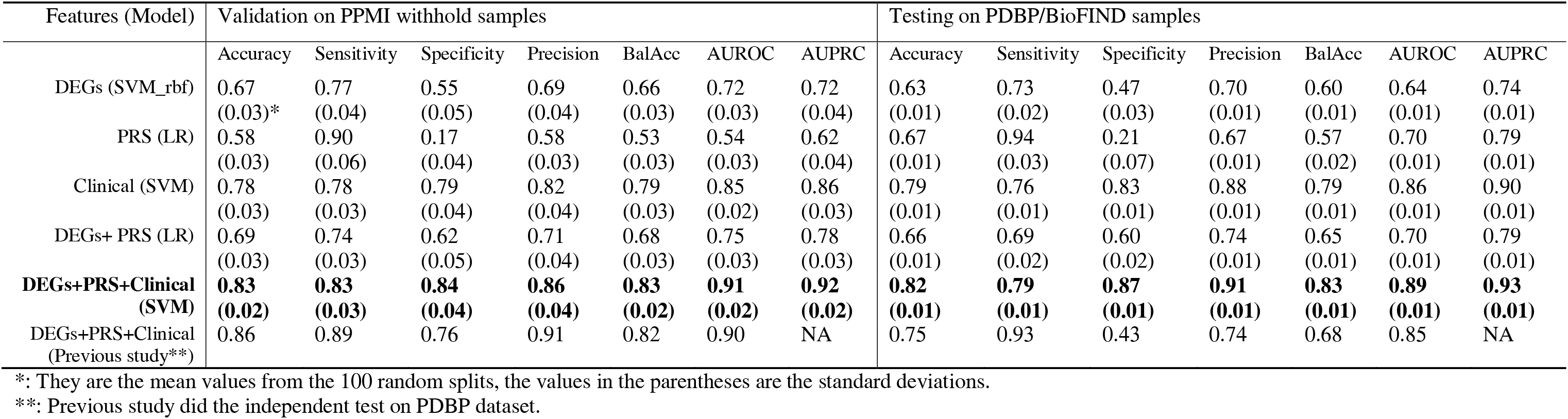
performance merits of classifier models on validation and testing datasets.

The comparisons of PD prediction potentials using DEGs, DE eRNAs, or DE circRNAs revealed that DE eRNAs models exhibit comparable performance to DE circRNAs models, but their predictive powers fall below those of DEGs models (**Fig S5, Supplementary Table S5**). Moreover, combining all DE eRNAs, DE circRNAs, and DEGs did not contribute to an enhancement in model performance. Further exploration into the PD prediction abilities of PRS or clinical data demonstrated that the best PRS-based model displayed similar AUROC and AUPRC values to the DEG+PRS model on the testing dataset (**Table 2, Supplementary Table S5)**. However, it exhibited lower precision, balanced accuracy values, and notably low specificity values, suggesting a tendency for PRS-based models to yield false positives. Additionally, PRS-based models had inferior performance on validation datasets compared to DEGs-based models. Therefore, DEGs proved effective in compensating for the shortcomings of PRS in predicting PD samples. Clinical data exhibited superior capabilities in distinguishing PD cases from healthy controls, although the performance values slightly lagged behind our final multi-omics model (**Table 2, Supplementary Table S5)**.

Delving into the selected features in each split, a total of 99 genes were chosen, with 12 of them recurrently selected as predictive features more than 80 times (**Fig 5F**, **Supplementary Table S5**). The remarkable consistency in feature selection, coupled with low standard deviations in performance values, affirmed the high stability of our model.

Looking into those selected genes, H19 is a long non-coding RNA, which was selected 100 times. H19 has been reported to be associated with PD progression and correlated with susceptibility to various CNS disorders(49, 50). We also found that 7 neutrophil genes were selected as the predictor during the 100 splits which includes *PREX1*, *SLCO4C1*, *CXCR2*, *DNAJC3*, *CD93*, *LAMP1*, *HEBP2*. The *LAMP1* were recurrently selected 98 times. *PREX1* and *CXCR2* were also the two genes that were replicated in brain data.

Above all, our final multi-omics model outperformed the recent publication with similar focuses respecting the accuracy (0.82 vs 0.75), balanced accuracy (0.83 vs 0.68), and AUROC (0.89 vs 0.85) on the testing dataset (**Table 2**). For the reasons why we had better performances than previous ripostes, it may because that we not only used the protein-coding genes as used in the previous study, but also included other genes in our DEGs as feature in our models. Additionally, we calculated the PRS using the 7,057 PD-associated significant variants instead of only the 90 SNPs that were used in previous published paper.

## Discussion

PD is a progressive, multisystem neurodegenerative disease that has been a huge burden on our society and the people it affects. Early diagnosis and biomarker discoveries that bolster the therapeutic pipeline for PD are urgently needed(51, 52). The Accelerating Medicine Partnership in Parkinson’s Disease (AMP PD) program has provided unprecedented opportunities for investigators, including this opportunity, to utilize the data to build an early diagnosis platform for PD patient diagnosis, which could lead to improved treatment response and higher efficacy. Currently, PD diagnosis is mainly based on clinical phenotype detections which could provide high sensitivity for detecting parkinsonism(15, 53). However, clinical observation alone is often insufficient to predict PD status prior to onset of disease. Once symptoms emerge and are detectable, it usually indicates developed PD status(10, 12). It has been reported that in idiopathic PD, there is severe degeneration of the nigrostriatal neurons of the substantia nigra before neurologists can establish the diagnosis according to the widely accepted clinical diagnostic criteria(54). It is conceivable that neuroprotective therapy starting at such a stage of the disease will fail to stop the degenerative process. Therefore, the identification of patients at risk and at earlier stages of the disease appears to be essential for any successful neuroprotection. The observational PD phenotypes are reflections of the changes in transcriptomic profiles, which were changing in advance of clinical phenotypes. Analyzing the transcriptomic changes between PD patients and healthy control samples can provide signals for preclinical diagnosis. Utilizing the large cohort datasets from AMP PD in this cross-sectional study, differentially expressed genes were initially discovered and then validated using these large sample size cohorts.

Functional enrichment analysis was conducted, and we found the neutrophil activation and degranulation were significantly enriched, which we would like to recommend serving as the diagnostic marker. As previously published, neutrophils infiltration plays important roles in the development of PD(55). Studies mentioned that circulating neutrophils are increased in numbers in PD while other circulating immune cells have decreased, or no change in prevalence(56, 57). In Craig *et al.* study utilizing the PPMI dataset found an increased number of neutrophils in PD patients compared to controls(23). While neutrophils have yet to be identified in the brains of PD patients, neutrophils have been identified in the brains of AD patients and mouse models of neuroinflammation(58, 59). Moreover, circulating neutrophils express *CD11b*, an integrin that responds to aggregated α-syn in microglia(60). Another study revealed that neutrophil degranulation was the most significantly altered molecular pathway in patients, with most genes in the neutrophil degranulation pathway containing nonsense or missense mutations(61). In our work we confirmed that the neutrophil activation and degranulation pathway were actively upregulated. By checking the publications and the pathway annotation databases(62, 63), we knew that neutrophils contain five different types of granules: primary granules, also known as azurophilic granules; secondary granules, also known as specific granules; tertiary granules; secretory vesicles and ficolin-rich granules. The primary granules are the main storage site of the most toxic mediators, including elastase, myeloperoxidase, cathepsins, and defensins. The secondary and tertiary granules contain lactoferrin and matrix metalloprotease 9 (also known as gelatinase B), respectively, among other substances. The secretory vesicles in human neutrophils contain human serum albumin, suggesting that they contain extracellular fluid that was derived from the endocytosis of the plasma membrane. Ficolin-rich granules are highly exocytosable gelatinase-poor granules found in neutrophils and rich in ficolin-1. Ficolin-1 is released from neutrophil granules by stimulation with fMLP or PMA. Granules are prevented from being released until receptors in the plasma membrane or phagosomal membrane signal to the cytoplasm to activate their movement to the cell membrane for secretion of their contents by degranulation. This is an important control mechanism as the neutrophil is highly enriched in tissue-destructive proteases.

There is increasing evidence showing the links between blood cells and PD development. Variants at or near the gene *LRRK2* locus have been known to be associated with PD. Reports shown that full-length *LRRK2* is a relatively common constituent of human peripheral blood mononuclear cells (PBMC) including affinity-isolated, CD14+ monocytes, CD19+ B-cells, and CD4+ as well as CD8+ T-cells(64). There was also evidence shown both *SNCA* mRNA and protein are particularly abundant erythroid cells(4). Lymphocyte is another category of cells that play important roles in PD. There are enhanced numbers of both CD4+ and CD8+ T cells in the brain parenchyma which had been observed in neuropathological studies of PD(65–67). A longitudinal case study of a PD patient found that alpha-synuclein-reactive T cells were most abundant in peripheral blood prior to the appearance of motor symptoms(68). Above all, more studies are emerging to show the potential diagnosis biomarkers in expression profiles of circulating genes.

TNF-alpha signaling pathway is another enriched pathway from our WikiPathway enrichment analysis. TNF-alpha had been approved to be increased both in the brain and in the cerebrospinal fluid from parkinsonian patients, and TNF-α is involved in the degenerative processes which occur in Parkinson’s disease. TNF-alpha is the key player in TNF-alpha signaling pathway, in our analysis, the leading-edge genes in this pathway includes *CFLAR, MAPK3, APAF1, PRKCZ, PYGL, MAP2K4, BTRC, NFKBIB, RAF1.* Currently, there are few studies having been focusing on these genes, so we my pay attentions to these gene in the future researches.

Previous studies have reported there is an association between the *SNCA* transcript abundance in blood with early stage and imaging-supported *de novo* PD. There is a paradoxical reduction in *SNCA* transcript counts in the blood of individuals with early-stage, neuroimaging-supported Parkinson’s disease(4, 31, 32). In our analysis, although the *SNCA* did not show the significant changes comparing the patient samples to health samples, we confirm the reduced trends in both our discovery and replication cohorts, as well as in our BRAINcode cohort. Besides, publication also showed that there is an inconclusive SNCA protein changes in plasma which is likely due to hemolysis of erythrocytes in which SNCA is one of the most plentiful proteins(4).

There have been some studies established the machine learning classifiers with different focuses and using different dataset. Scherzer *et.al* built the first ML classifier in PD using 22 genes. Liu *et.al* used clinical and genetic information for the prediction of cognitive decline in patients with Parkinson’s disease and the progression of PD(69, 70), and Severson *et.al* identified subtypes of PD based on clinical data(71). Here, to maximize the value of the massive amount of data, we tested several machine-learning methods for PD diagnosis classification make use of the clinical data, transcriptomics data and genetics data. Our final multi-omics model has high AUC values and high sensitivity and specificity, which means our model cannot only tell the true PD patients but also recognize the low-risk people. For the next step, we may try more advanced machine algorithms, such as the DNN, CNN, and VAE, to improve the performances and dig for more meaningful insights behind the data.

We may still have a few limitations of the current analysis. Our current analysis was focused on the diagnostic classification of PD at the baseline in a cross-sectional design. Future analyses will be important to prospectively and longitudinally test diagnostic classifiers. Moreover, progression markers are needed, and this will require analyses of longitudinal RNA data sets. In order to begin to translate these candidate classifiers to the clinic, more research is needed to clarify the positive and negative predictive values in different clinically relevant scenarios, e.g., as aid for augmented medicine in the movement disorders clinic populations or as screening tool for high-risk individuals in the general population. These scenarios involve highly distinct incidence of PD patients and require a clearer understanding of positive predictive value (PPV) and negative predictive value (NPV).

Overall, in this study, we identified a set of DE RNAs and defined neutrophil activation and degranulation as potential early diagnostic biomarkers. We built a high-performance PD classification model which could be helpful for PD diagnosis prediction. We provided a computational framework that will be helpful for PD biomarker discovery and provide disease risk prediction, which is a critical step for better assessment of PD risk and accelerating the PD disease diagnosis.

## Methods

### Study design

First, we discovered genes that are differentially expressed in PD in an analysis of the discovery cohort and replicated those significant genes in a cross-sectional analysis of the replication cohort. In our previous study, we probed the transcriptome of dopamine neurons in post-mortem brains with various levels of neuropathology. We then evaluated the blood-based PD-associated genes (discovered and replicated DEGs) for association with PD neuropathology in dopamine neurons using our laser-captured RNA-seq dataset (BRAINcode, http://www.humanbraincode.org)(6). Meanwhile, the functional enrichment analysis was conducted on those replicated DEGs. As well, the cell type enrichment analysis was carried out to find the enriched cell types of the replicated DEGs. Furthermore, the eRNAs and circRNAs were quantified in both discovery and replication datasets and the replicated significantly differentially expressed eRNAs (DE eRNAs) and circRNAs (DE circRNAs) were presented in this work. Utilizing the DEGs, genetics and clinical data, we built the PD diagnosis classifier models. (**Fig 1**).

### Sample and gene expression quality control

Filters were applied to remove those participants as shown in **Fig S2** and **Fig S3**. The same filtration strategies were applied to both the discovery and the replication datasets. At the very beginning, participants without RNA-seq were removed. In the next step, only the patients that have the baseline RNA-seq data with RIN greater than 5.0 were kept in our following analysis. To limit batch effects due to ancestry, we restricted our analysis to patients self-identifying as White. Meanwhile, we restricted our analysis to patients listed as either cases or controls. Lastly, we excluded those participants with diagnosis conflicts during the follow-up visits after the initial enrollment in case and control groups separately. Those PD cases whose diagnosis changed during follow up were removed. Similarly, control participants who developed PD were excluded. Prodromal participants and SWEDD (Scans without evidence of dopaminergic deficit) patients were also removed. Participants with missing clinical or genetic data were also moved as those data would be used in the following analysis.

Quality control of expression data was performed to filter out lowly expressed genes and remove samples outliers. For the genes, we first removed genes that have low expression levels defined as counts fewer than 5 reads in more than 90% of samples and variances of less than 1 across the samples. To check if there is any sex information that is mislabeled, a scatter plot of the expression levels of a Y chromosome-specific gene and an X chromosome-specific gene was plotted. We also verified the biases of sequencing data arising from case/control sample distributions on the plates were minimal.

### Identification of PD-associated mRNAs

The differential expression analysis was conducted using DEseq2 (v1.36.0)(72). The gene read counts data from Salmon(73) quantification result files was used. The primary differential expression was tested between the PD conditions (PD cases vs. healthy controls), and the age_at_baseline (continuous variable), sex, plate, RIN, and the top 10 principal components (PCs) of the genotype data were included as covariates in DEseq2. The replicated DEGs were further analyzed using ClusterProfiler(74) to find the enriched functions. We also performed cell-type-specific enrichment analysis using the WebCSEA online tool(75) to find which human tissue-cell types these genes might manifest their impacts on.

As a secondary analysis, we further looked at gene expression changes associated with motor severity, indicated by the MDS-UPDRS part III summary score. Tests were performed in the same DESeq2 framework where the MDS-UPDRS score was treated as a continuous dependent variable.

### Identification of PD-associated enhancer RNAs and circular RNAs

Since the AMP PD provided the raw whole sequencing data, we would like to know the non-coding novel RNAs, especially the eRNA and circRNA differences in PD patients and healthy individuals. We called eRNAs and circRNAs in all datasets. We used our previously developed method(6) to identify eRNA candidates in the blood. The circRNAs were called using the CIRCexplorer2 package(76). Then differential expression analysis was conducted on the eRNA and circRNA reads count using DESeq2. Since the circRNAs have relatively lower reads count in the samples, we used all samples, instead of the baseline samples only, to increase the sample size in order to empower the DE circRNA discovery. The same covariates as in finding the DEGs were used.

### Confirmation in brain

We tested blood biomarkers transcripts using the BRAINcode dataset. The PD-associated RNAs that are also differentially expressed in brain will be highly relevant and prioritized for validation. We conducted the DE analysis using the data from brain neuron samples and compared the blood DEGs and brain DEGs. In our BRAINcode v2 project, we performed laser-capture microdissection total RNA-sequencing (lcRNAseq)(3) on dopamine neurons from the midbrain substantia nigra pars compacta of 104 high-quality human postmortem brains (HC: n = 59; ILB: n = 27; PD: n=18). Many polyadenylated and non-polyadenylated transcripts are identified with high confidence. The DEGs in dopamine neurons were identified between PD samples and health control samples. The DEGs with the same fold change directions as in brain data were obtained.

### Evaluation on a second digital gene expression platform in the Harvard Biomarkers Study

We also compared the expression levels of several PD-associated genes from the blood with our in-house NanoString data to validate our findings. The NanoString dataset with PD cases and healthy controls are nested in the Harvard Biomarker Study (HBS). The participants’ blood sample with high RNA quality (RIN ≥ 7) was processed for digital expression analysis on the NanoString platform(77) with 33 distinct molecular barcodes (29 PD-associated genes) to count the abundance of selected-transcripts directly in RNA from blood cells. A total of 617 PD cases and 618 healthy controls passed normalization processing for validating our findings.

### Construction of PD diagnosis classifier models

We have built the classifiers utilizing the multi-modality data which includes transcriptomics, polygenic risk score (PRS), and clinical data. The PPMI samples (discovery cohort) were randomly split into a training set (80%) and a validation set (20%). We did the randomly splits 100 times to test the model stability. The training set was used to build the classifiers. The validation set was used to optimize the hyper-parameters of each model through a grid search. Final models were tested on the independent PDBP/BioFIND samples (replication cohort).

Three models were built in sequential order using the following feature sets respectively: transcriptomics only (“DEGs”), transcriptomics plus polygenic risk score (“DEGs+PRS”), and transcriptomics, polygenic risk score, and clinical data combined (“DEGs+PRS+Clinical”). The transcriptomics data is the 874 DEGs from the PPMI cohort. The PRS was calculated using PRSice-2(78) based on the 7,057 PD-associated significant variants from the recently published PD GWAS work(14). Clinical data includes the total UPSIT score, sex, and age at the baseline. Since we have too many features, and the feature size is larger than the sample size, we have tried feature selections and modeling the classifiers to avoid overfitting.

To train the models, the variance stabilizing transformed (VST) expression abundances were standardized after log transformations. Feature selection was conducted on the training set using the LASSO approach by making use of sklearn.linear_model.Lasso function and the parameter alpha were screened to pick the best one in order to have the best area under the receiver operating characteristic curve (AUROC) value. Only features with non-zero coefficients were included in the model. To take the advantages of different machine learning algorithms, 10 different machine learning classifiers were trained and compared, including support vector machine with linear kernel (SVM), support vector machine with rbf kernel (SVM_rbf), linear regression (LN), logistic regression (LR), stochastic gradient descent (SGD), AdaBoost classifier (ABC), gradient boosting classifier (GBC), random forest (RF), k-nearest neighbors (KNN), and multiple layers perceptron classifier (MLP).

To investigate if the eRNAs, or circRNAs are predictive for PD diagnosis, we also tested the classifiers using the DE eRNAs, and DE circRNAs separately.

## Data Availability

The PPMI, PDBP, and BioFIND data can be accessed from the AMP-PD Google Cloud Storage with the approved “Data Use Approvement”. All the up-to-date information and data collection or data processing procedures on the AMP-PD program can be found at https://www.amp-pd.org. The brain neuron data and the NanoString were produced in our own lab. All the analysis code can be accessed at: https://github.com/bwh-bioinformatics-hub/AI2AMP-PD.

## Acknowledgment

This study was funded in part by NIH grant 1U01NS120637, R01AG057331, U01 NS082157, the U.S. Department of Defense (to C.R.S.), the American Parkinson Disease Association (APDA) Research Award (to X.D.). C.R.S.’s work is supported by NIH grants NINDS/NIA R01NS115144, U01NS095736, U01NS100603, and the American Parkinson Disease Association Center for Advanced Parkinson Research. X.D. received funding from the American Parkinson Disease Association (APDA). C.R.S and X.D.’s work was in part funded by the joint efforts of The Michael J. Fox Foundation for Parkinson’s Research (MJFF) and the Aligning Science Across Parkinson’s (ASAP) initiative. MJFF administers the grant [ASAP-000301] on behalf of ASAP and itself. Data used in the preparation of this article were obtained from the Accelerating Medicine Partnership® (AMP®) Parkinson’s Disease (AMP PD) Knowledge Platform. For up-to-date information on the study, visit https://www.amp-pd.org. For the purpose of open access, the author has applied a CC BY public copyright license to all Author Accepted Manuscripts arising from this submission.

---

**Fig S1.**
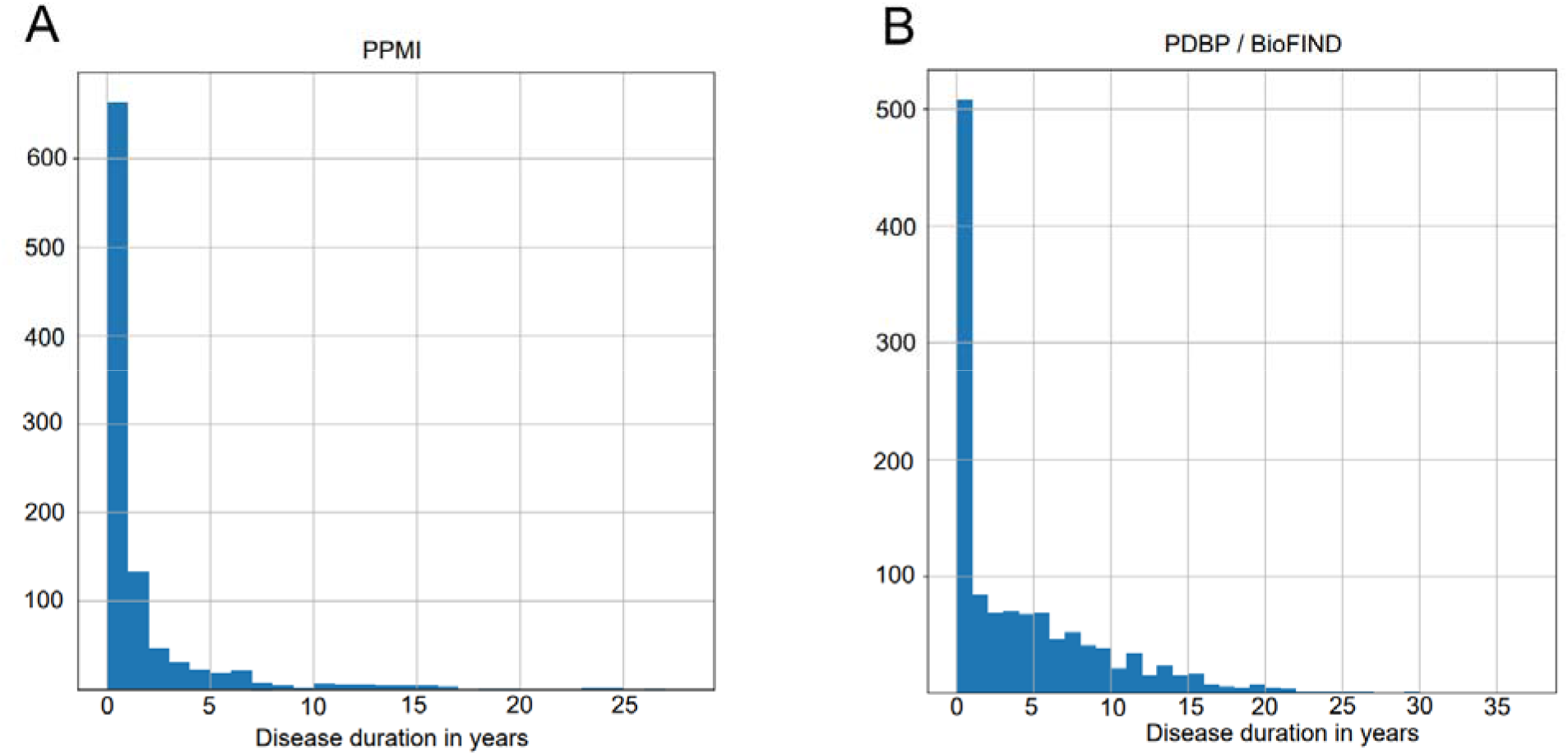
The distributions of disease duration at enrollment in discovery (A) and replication (B) cohort.

**Fig S2.**
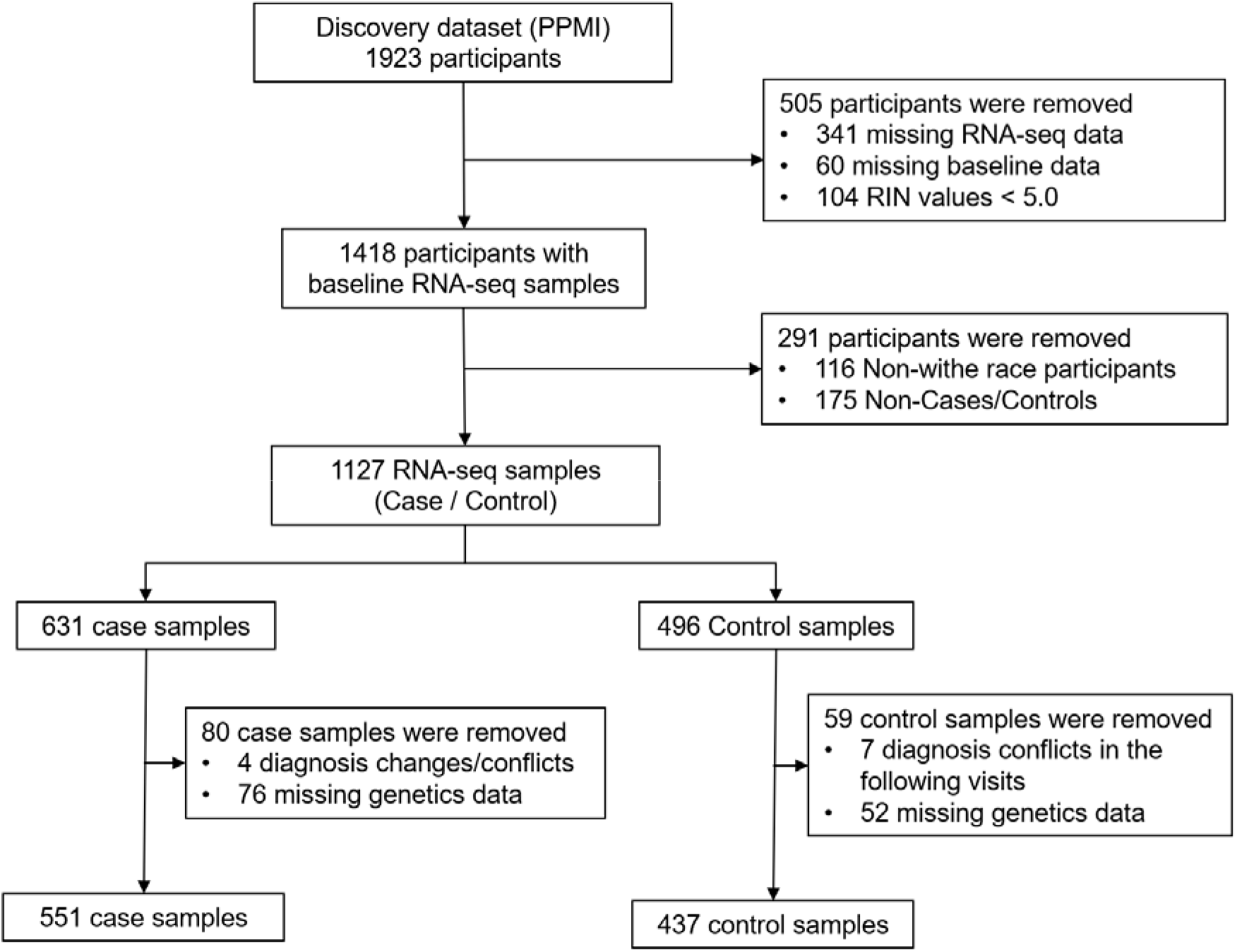
The steps on the discovery dataset.

**Fig S3.**
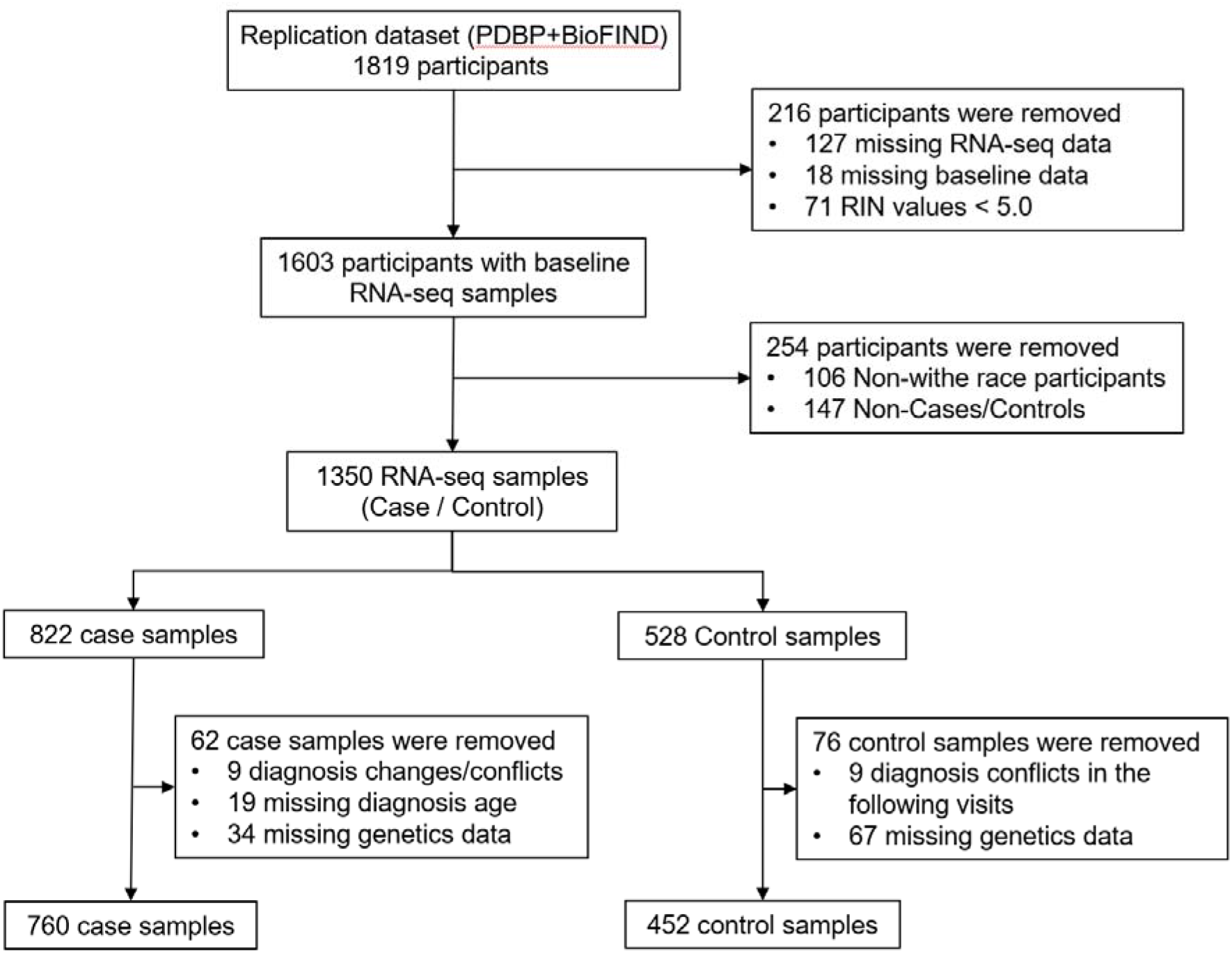
The steps on the replication dataset.

**Fig S4.**
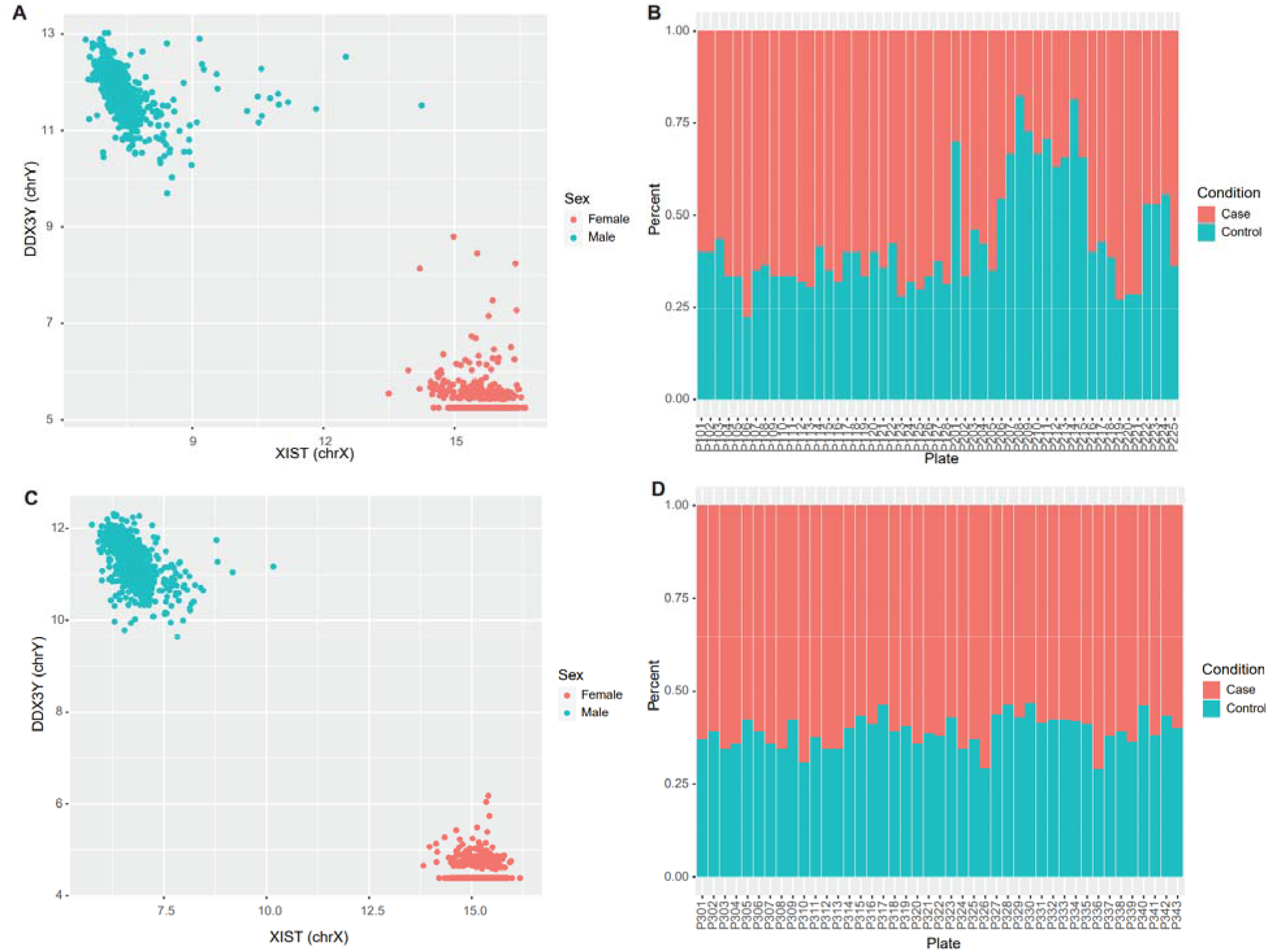
Data quality control assessments in the discovery and replication datasets. (A, B) Sex check and the samples distributions on the plate of discovery dataset. (C, D) Sex check and the samples distributions on the plate of replication dataset.

**Fig S5.**
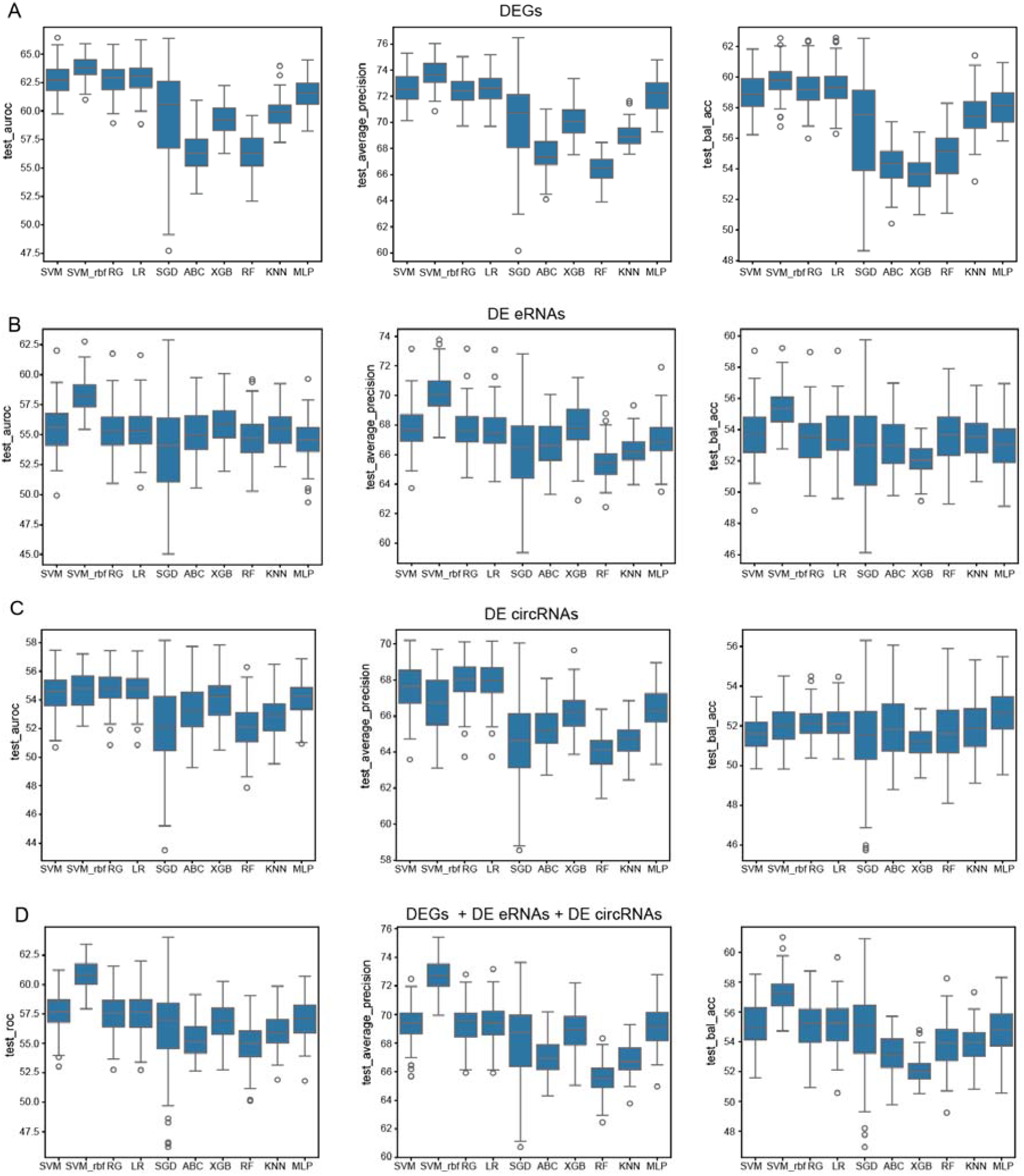
The AUROC, average precision and balanced accuracy values of the models using DEG, DE eRNAs, or DE circRNAs as features.

