## Supplementary material for "Transcriptional pathobiology and multi-omics predictors for Parkinson’s disease": Table 1

Table 1. Summary of clinical data at baseline (after filtrations.)

|  |  | Discovery dataset | | Replication dataset | | p-value |
| --- | --- | --- | --- | --- | --- | --- |
|  |  | PD (n=551) | HC (n=437) | PD (n=760) | HC (n=452) |  |
| Sex | Male | 318 | 214 | 482 | 204 | 1.59*10^-4^ |
|  | Female | 233 | 223 | 278 | 248 | 0.6086 |
| Age, years (mean(sd)) | | 62.91 (10.42) | 57.18 (12.53) | 64.99 (8.78) | 62.94 (10.81) | 0.8984 |
| Duration of disease (mean(sd)) | | 2.37 (4.12) | \ | 6.19 (5.18) | \ | 5.98*10^-41^ |
| RIN value (mean(sd)) | | 8.18 (0.90) | 8.36 (0.84) | 7.44 (0.83) | 7.41 (0.85) | 1.0 |
| MOCA (mean(sd)) | | 24.86 (4.17) | 27.94 (1.88) | 25.30 (3.60) | 26.53 (2.54) | 1.0 |
| UPSIT (mean(sd)) | | 22.22 (8.03) | 33.6 (4.70) | 19.62 (7.74) | 32.48 (6.00) | 0.8429 |
