## Supplementary material for "Transcriptional pathobiology and multi-omics predictors for Parkinson’s disease": Table 2

Table 2. performance merits of classifier models on validation and testing datasets

| Feature dataset (Model) | Validation on PPMI withhold samples | | | | | | | Testing on PDBP/BioFIND samples | | | | | | |
| --- | --- | --- | --- | --- | --- | --- | --- | --- | --- | --- | --- | --- | --- | --- |
|  | Accuracy | Sensitivity | Specificity | Precision | BalAcc | AUROC | AUPRC | Accuracy | Sensitivity | Specificity | Precision | BalAcc | AUROC | AUPRC |
| DEGs (SVM_rbf) | 0.67 (0.03)* | 0.77 (0.04) | 0.55 (0.05) | 0.69 (0.04) | 0.66 (0.03) | 0.72 (0.03) | 0.72 (0.04) | 0.63 (0.01) | 0.73 (0.02) | 0.47 (0.03) | 0.70 (0.01) | 0.60 (0.01) | 0.64 (0.01) | 0.74 (0.01) |
| PRS (LR) | 0.58 (0.03) | 0.90 (0.06) | 0.17 (0.04) | 0.58 (0.03) | 0.53 (0.03) | 0.54 (0.03) | 0.62 (0.04) | 0.67 (0.01) | 0.94 (0.03) | 0.21 (0.07) | 0.67 (0.01) | 0.57 (0.02) | 0.70 (0.01) | 0.79 (0.01) |
| Clinical (SVM) | 0.78 (0.03) | 0.78 (0.03) | 0.79 (0.04) | 0.82 (0.04) | 0.79 (0.03) | 0.85 (0.02) | 0.86 (0.03) | 0.79 (0.01) | 0.76 (0.01) | 0.83 (0.01) | 0.88 (0.01) | 0.79 (0.01) | 0.86 (0.01) | 0.90 (0.01) |
| DEGs+ PRS (LR) | 0.69 (0.03) | 0.74 (0.03) | 0.62 (0.05) | 0.71 (0.04) | 0.68 (0.03) | 0.75 (0.03) | 0.78 (0.03) | 0.66 (0.01) | 0.69 (0.02) | 0.60 (0.02) | 0.74 (0.01) | 0.65 (0.01) | 0.70 (0.01) | 0.79 (0.01) |
| **DEGs+PRS+Clinical (SVM)** | **0.83 (0.02)** | **0.83 (0.03)** | **0.84 (0.04)** | **0.86 (0.04)** | **0.83 (0.02)** | **0.91 (0.02)** | **0.92 (0.02)** | **0.82 (0.01)** | **0.79 (0.01)** | **0.87 (0.01)** | **0.91 (0.01)** | **0.83 (0.01)** | **0.89 (0.01)** | **0.93 (0.01)** |
| DEGs+PRS+Clinical (Previous study**) | 0.86 | 0.89 | 0.76 | 0.91 | 0.82 | 0.90 | NA | 0.75 | 0.93 | 0.43 | 0.74 | 0.68 | 0.85 | NA |

*: They are the mean values from the 100 random splits, the values in the parentheses are the standard deviations.

**: Previous study did the independent test on PDBP dataset.
